# Extension of the *in vivo* haploid induction system from maize to wheat

**DOI:** 10.1101/609305

**Authors:** Chenxu Liu, Yu Zhong, Xiaolong Qi, Ming Chen, Zongkai Liu, Chen Chen, Xiaolong Tian, Jinlong Li, Yanyan Jiao, Dong wang, Yuwen Wang, Mengran Li, Mingming Xin, Wenxin Liu, Weiwei Jin, Shaojiang Chen

## Abstract

Doubled haploid breeding technology has been one of the most important techniques for accelerating crop breeding. In compare to *in vivo* haploid induction in maize, which is efficient and background independent, wheat haploid production by interspecific hybridization pollinated with maize is influenced by genetic background and requires rescue of young embryos. Here, we analyzed the homologues of maize haploid induction gene *MTL*/*ZmPLA1*/*NLD* in several crop species systematically, the homologues are highly conserved in sorghum, millet and wheat etc. Since wheat is a very important polyploidy crop, as a proof of concept, we demonstrated that the *in vivo* haploid induction method could be extended from diploid maize to hexaploid wheat by knocking out the wheat homologues (*TaPLAs*). Result showed that double knock-out mutation could trigger wheat haploid induction at ~ 2%-3%, accompanied by 30% - 60% seed setting rate. The performance of haploid wheat individual showed shorter plant, narrower leaves and male sterile. Our results also revealed that knockout of *TaPLA*-A and *TaPLA*-D do not affect pollen viability. This study not only confirmed the function of the induction gene and explored a new approach for haploid production in wheat, but also provided an example that the *in vivo* haploid induction could be applied in more crop species with different ploidy levels. Furthermore, by combining with gene editing, it would be a fast and powerful platform for traits improvement in polyploidy crops breeding.

## Introduction

Doubled haploid (DH) technology substantially accelerates the breeding process for several crop species. Several different methods have been established for producing haploids in different crop species (Ishii et al., 2016), including interspecific hybridization between maize and wheat to produce wheat haploids (Laurie, 1988); *in vivo* haploid induction to produce maize haploids (E. H. Coe, 1959), and anther/male gametophyte culture method to produce haploids in rice and other crop species (Germanà, 2011). Of these, only *in vivo* haploid induction in maize has been demonstrated to be independent of genetic background and to produce haploids efficiently (> 10%), which make it possible to produce DH lines in an engineered way and laying a solid foundation for modern commercial breeding (Chen, 2013; Geiger, 2009). Recent studies of the genetics underlying *in vivo* haploid induction in maize revealed that several quantitative trait loci (QTL) contributed to haploid induction rate (HIR) (Prigge et al., 2012). Of these, *qhir1* is critical for haploid induction (Dong et al., 2013; Liu et al., 2015). Loss of function of the gene *MTL*/*ZmPLA1*/*NLD* in *qhir1* triggers haploid induction (Gilles et al., 2017; Kelliher et al., 2017; Liu et al., 2017a). Importantly, *in vivo* haploid induction system had been successfully applied to rice (Yao et al., 2018), making this method more important in plant breeding. Genetic analysis on the QTLs contributing HIR revealed that *qhir8* can improve haploid induction efficiency by 3- to 5- fold (Liu et al., 2015), this is a promising result for improving DH breeding efficiency in crop species other than maize. In addition to producing homozygous DH lines, *in vivo* haploid induction in maize has also been used for gene editing without introducing the genome of the male parent, which is meaningful for genome editing in different genetic backgrounds (Kelliher et al., 2019; Wang et al., 2019).

Compared with *in vivo* haploid induction in maize, wheat haploid production through interspecific hybridization requires the rescue of young embryos and is depend on genetic background (Niu et al., 2014), which limits its application. *In vivo* haploid induction system has been extended from maize to rice, while both are diploid crop, little is known whether it can be applied to polyploidy plants. Wheat is a very important hexaploidy crop, to extend maize *in vivo* haploid induction to wheat would create a novel protocol in speeding up wheat breeding and providing a potential method for gene editing in different genetic backgrounds (Kelliher et al., 2019; Wang et al., 2019). Furthermore, it would also give an example for exploring the haploid production and editing system in other polyploidy crops.

## Results and Discussion

The full length amino acid sequence encoded by *MTL/ZmPLA1/NLD* was used to do a BLAST search for homologues in different crop species (www.gramene.com), followed by amino acid alignment and phylogenic analysis using MEGA (Knyaz et al., 2018). Results showed that the gene is highly conserved among 19 species of Liliopsida (Supplemental Figure 1). Sorghum and millet homologues had >80% amino acid sequence identity with maize, and wheat and rice homologues were divided into two different branches (Supplemental Figure. 1), both which sharing ~70%-80% amino acid sequence identity with maize. Wheat genome consists three homologues: TraesCS4A02G018100 (*TaPLA*-A), TraesCS4B02G286000 (*TaPLA*-B) and TraesCS4D02G284700 (*TaPLA*-D), located on chromosomes 4A, 4B and 4D, respectively. All three shared a similar gene structure that includes four exons (Figure1. A). The amino acid sequence identity between each of the three homologues and *MTL*/*ZmPLA1*/*NLD* is 75%, representing a high level of conservation property. The amino acid sequences identity among the three *TaPLA* genes is 96%.

**Figure 1.**
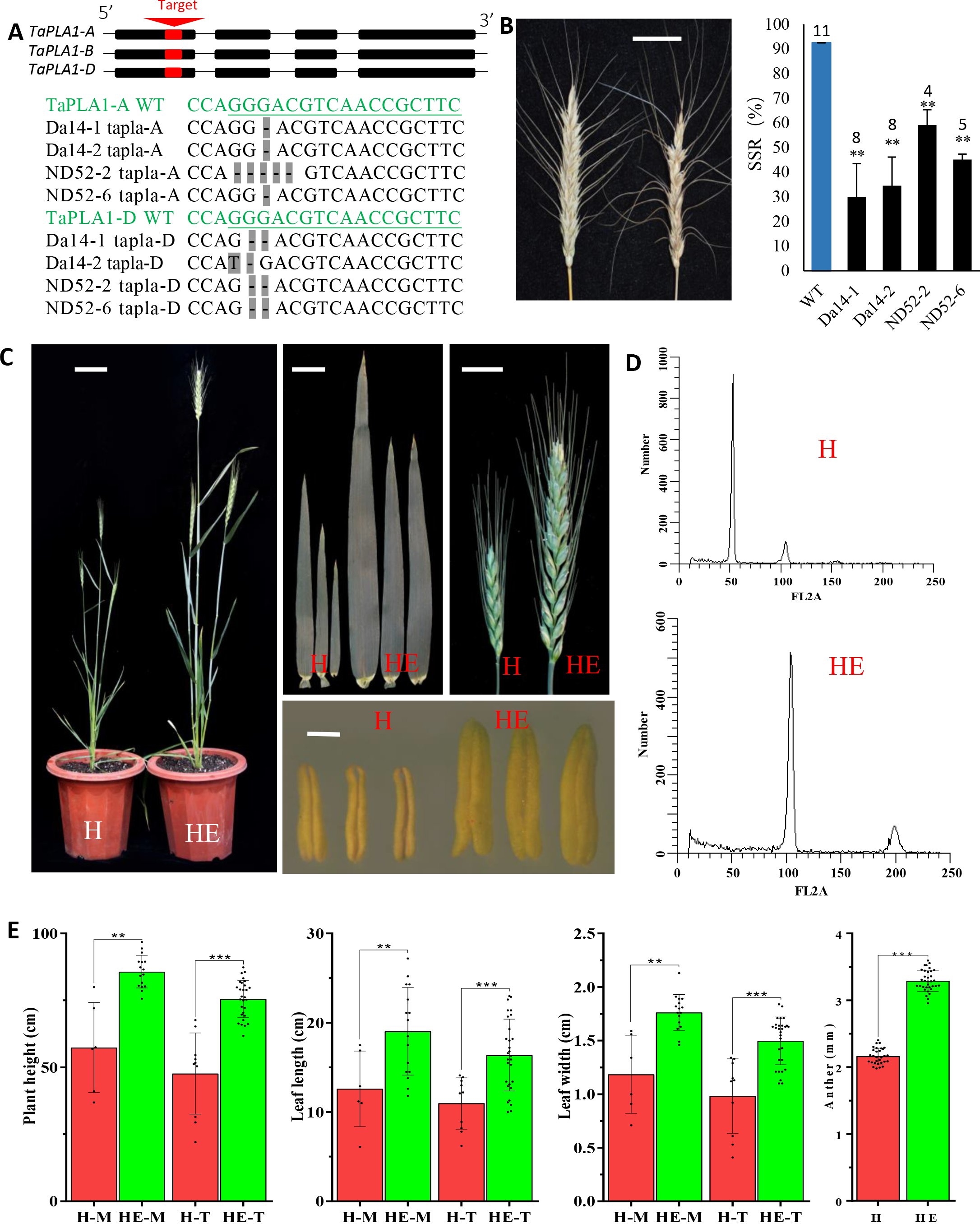
Knockout of wheat homologues of *MTL*/*ZmPLA1*/*NLD* and phenotypic analysis. **(A)** Gene structure of wheat homologues *TaPLA*-A, *TaPLA*-B and *TaPLA*-D. Exons in red were target regions of guide RNAs. Shown below are knockout sequence results for of transgenic line Da14-1, Da14-2, ND52-2 and ND52-2 at *TaPLA*-A and *TaPLA*-D, respectively. Sequences in green are target sequences of *TaPLA*-A and *TaPLA*-D, the guide RNA is underlined. Black indicates the sequence of transgenic events. Deletions are shown by “-”; on gray background. **(B)** Ear performance of in wildtype (left) and knockout (right) plants. Statistical analysis for seed setting rate (SSR) of wildtype and four transgenic lines are shown on the right side. SSR was calculated as SSR (%) = (number of viable seeds/number of embryo sacs on an ear) ×100%. ** P < 0.01 calculated with the heteroscedastic two-tailed Studentx2019;s *t-*test. Sample size of SSR for wildtype and each transgenic line was marked on the top of column. **(C)** Phenotypic difference between haploid (H) and hexaploidy (HE) wheat, including whole plant, leaf, ear and anther. H, haploid; HE, Hexaploid. Bars = 5 cm, 1 cm, 1 cm and 1 mm for whole plant, flag leaf, ear and anther, respectively. **(D)** Ploidy verification by flow cytometry for hexaploid and haploid plants. **(E)** Statistical analysis of phenotypic differences including plant height, leaf length, leaf width and anther length between haploid and hexaploid wheat. Both stems (M) and tillers (T) of haploids and hexaploids were analyzed. A heteroscedastic two-tailed Student’s *t*-test was performed for each class between wildtype and mutant. *, ** and *** P<0.05, 0.01 and 0.001, respectively.

Next, we designed a guide RNA sequence (Supplemental Table 1) targeting the first exon of all three genes to create complete knockout lines using double-target CRISPR/Cas9 system (Figure1. A) (Xing et al., 2014). After *Agrobacterium tumefaciens*-mediated transformation, four transgenic events were obtained with mutations in both *TaPLA*-A and *TaPLA*-D. None of them had a mutation in *TaPLA*-B in subgenome B. Of the four transgenic events obtained, two transgenic lines, Da14-1 and ND52-6, were found to carry the same mutations: a 1bp guanine (G) deletion in *TaPLA*-A and a GG deletion in *TaPLA*-D. Da14-2 had the same 1-bp deletion in TaPLA-A as Da14-1 and ND52-6, but had a G to cytosine (C) mutation and a 1-bp G deletion in the gene *TaPLA*-D (Figure1. A). ND52-2 had a 5-bp GGGAC deletion in *TaPLA*-A and a GG deletion in *TaPLA*-D. Although different in sequence, all these mutations led to frame shifts and loss of function for both *TaPLA*-A and *TaPLA*-D (Figure1. A). These transgenic were chosen for subsequent analysis.

All T_0_ transgenic and control plants were grown in the same greenhouse. No substantive phenotypic difference was found between wildtype and transgenic plants (Supplemental Table 1), except for the seed setting rate (SSR), which ranged from ~30% to ~60% in transgenic lines, significantly lower than the average value of 92.6% in wildtype (Figure1. B), implying that *TaPLAs* may be involved in sexual reproduction. Haploid induction in transgenic lines was evaluated in the self-pollination progenies of T_0_ generation. Putative haploid wheat plants were identified according to the haploid plant characteristic reported for maize and wheat (Liu et al., 2017a; Zhang et al., 2014). Putative haploid plants were found in self-pollinated progenies of all four transgenic lines (Table 1). In Da14-1 and Da14-2, two putative haploids were found among their T_1_ populations of 70 and 82 plants, respectively. In ND52-2, three putative haploids occurred in a T_1_ population of 83 plants. In ND52-6, four putative haloids were identified among 102 plants (Figure1. C, Table 1). Flow cytometry was used to determine the real ploidy of each putative haploid. Compared with hexaploid controls which had an FL2A peak approximately 100, all putative haploids had an FL2A approximately 50 (Figure1. D), suggesting that putative haploids identified by phenotypic characteristic were true haploids. Compared with the hexaploid wheat plants, these haploids had shorter plant height, narrower leaves, shorter ears and male sterility (Figure1. C, E). No haploid plant was identified in a control group with 267 wildtype individuals. Therefore, we conclude that knockout of wheat homologues of *MTL*/*ZmPLA1*/*NLD* could trigger wheat haploids induction.

**Table 1.** Haploid induction potential of transgenic lines. Haploid plants occurred in self-pollination progeny of transgenic lines, genotype on *TaPLA*-A and genotype *TaPLA*-D showed, H, heterozygous; M, homozygous mutant; W, wildtype. The control group comprised self-pollinated plants from the wildtype receptor background CB037.

| Transgenic<br>events&Control | Genotype<br>*TaPLA-A* | Genotype<br>*TaPLA-D* | Total<br>plants | Haploid<br>plants | HIR (%) |
| --- | --- | --- | --- | --- | --- |
| Da14-1 | M | H | 70 | 2 | 2.86 |
| Da14-2 | H | H | 82 | 2 | 2.44 |
| ND52-2 | H | H | 83 | 3 | 3.61 |
| ND52-6 | H | H | 102 | 4 | 3.92 |
| Control | W | W | 267 | 0 | 0.00 |

To characterize the expression pattern of the *TaPLA* genes, subcellular localization was performed in tobacco epidermal cells infected by agrobacterium. As shown in Supplemental Figure. 2, all three genes showed strong signals in the cytomembrane, that merged well with the endoplasmic reticulum (ER) marker. This result is consistent with the cytomembrane subcellular localization of *MTL*/*ZmPLA1*/*NLD* in maize (Gilles et al., 2017). Reciprocal crosses between the maize haploid inducer and non-inducer lines previously revealed that the transmission of male gametes from inducer lines is decreased significantly (Dong et al., 2013; Liu et al., 2017a), suggesting that haploid inducer lines lead to either low pollen viability of pollen or a defect in fusion between male and female gametes in double fertilization. Therefore, we performed fluorescein diacetate (FDA) staining to examine pollen viability in wildtype and mutants. Pollen from wildtype and mutant lines was divided into three classes: high-viability pollen, low-viability pollen and dead pollen according to their pollen viability (Figure2. A). There was no difference in the proportion of pollen viability classes between mutants and wildtype (Figure2. B). This result demonstrates that loss of function of both *TaPLA*-A and *TaPLA*-D does not influence pollen viability.

**Figure 2.**
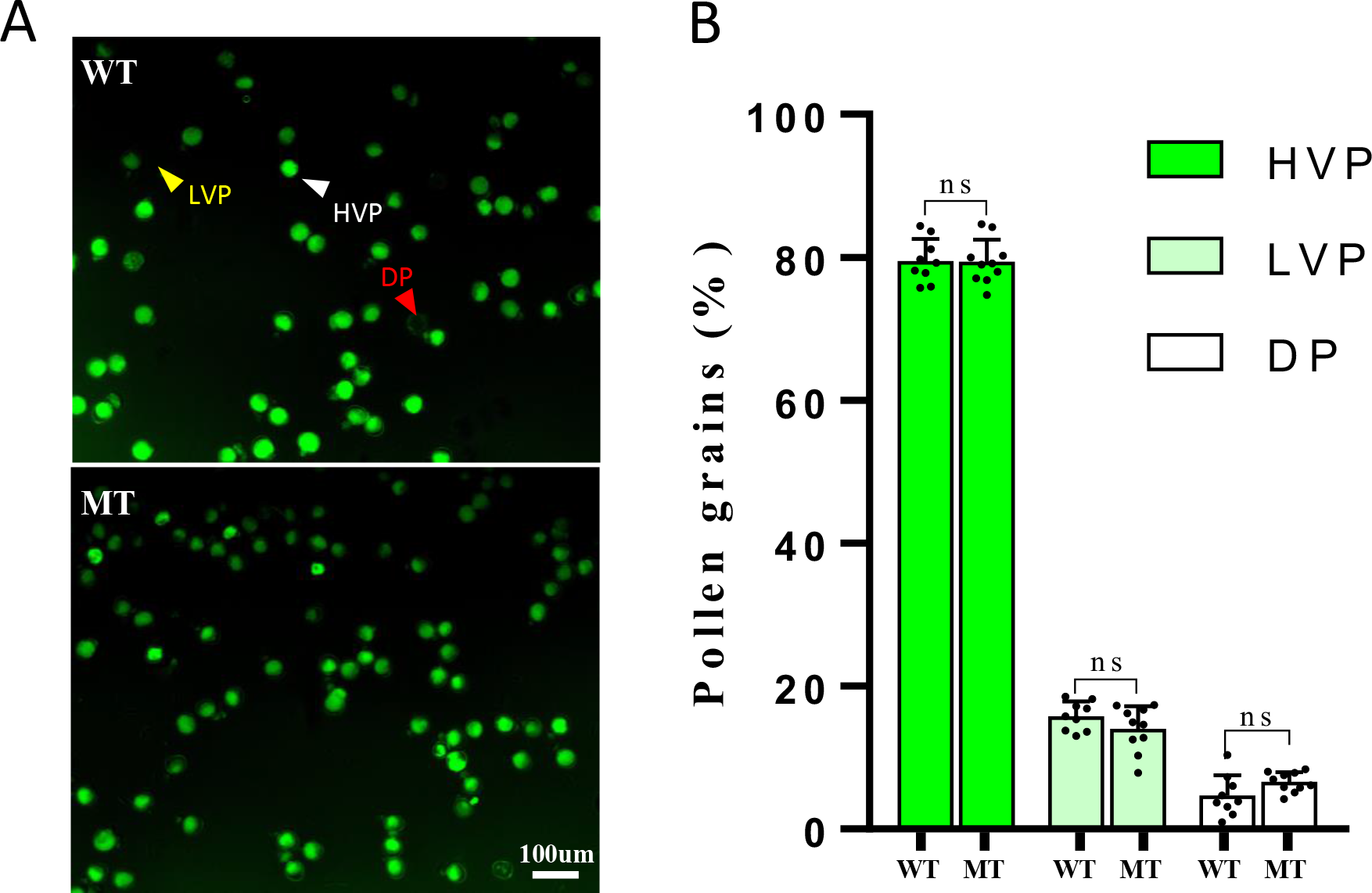
Pollen viability assays with FDA staining. **(A)** Comparison of pollen viability between wildtype and mutants in CB037 background (scale bar, 100μ). Pollen with a strong GFP (white triangle) and weak (yellow triangle) or no (red triangle) represented for high-viability pollen (HVP), low-viability pollen (LVP) and died pollen (DP), respectively. **(B)** Ratio of the three viability classes in wildtype and mutant pollen. Statistics were conducted with three biological replicates. A heteroscedastic two-tailed Studentx2019;s *t*-test was performed for each class between wildtype and mutant. NS, not significant P≥ 0.05).

In summary, as a proof of concept, we have demonstrated that *in vivo* haploid induction can be extended from diploid maize to wheat, by knocking out *TaPLA*-A and *TaPLA*-D. Previous studies on *in vivo* haploid induction in crop species, used maize or rice, which are diploid. This study provided the first proof that *in vivo* haploid induction is not limited to diploid crop species but can be extended to polyploidy species, which substantially extended its potential. Considering the potential redundancy among *TaPLA*-A, *TaPLA*-B and *TaPLA*-D, existence of *TaPLA*-B may have functional complementation effect in *Tapla*-A and *Tapla*-B double mutant. Nevertheless, the complementation effect is not sufficient to rescue the phenotype of haploid induction and decreased SSR. On the other hand, further improvement in the efficiency of haploid induction in wheat may be achieved by creating triple mutants.

The observed phenotype of *TaPLAs* knockout lines including a ~2-3% haploid frequency and significantly decreased SSR, are similar to that of maize and rice (Gilles et al., 2017; Kelliher et al., 2017; Liu et al., 2017a; Yao et al., 2018), suggesting that *MTL*/Zm*PLA1*/*NLD* homologues are involved in the similar pathway during double-fertilization across different crop species. Thus, our study provided an alternative crop species in exploring the biological mechanism of haploid induction.

Several technical problems require further investigation before *in vivo* haploid induction can be applied efficiently in wheat, including low efficiency in haploid induction and the difficulties in wheat haploid identification in seed stage. Nonetheless, our study provides an alternative approach other than interspecific hybridization for generating haploids in wheat. Through in-depth study of the genetic basis and biological mechanisms of *in vivo* haploid induction in maize, more and more genes contributing HIR will be identified (Liu et al., 2015; Prigge et al., 2012). The efficiency of wheat haploid induction may be further improved by the knockout/over-expression of these new identified homologous genes contributing HIR in maize. On the other hand, recent studies enable the identification of haploids using enhanced green fluorescent proteins (EGFP) and DsRED signals specifically expressed in the embryo and endosperm, respectively (Dong et al., 2018). This method may provide a potential solution for haploid kernel identification in wheat, which lacks phenotypic markers like *R1-nj* and high oil content existed in maize (Dong et al., 2014).

Recent studies that utilized the combination of CRISPR/Cas9-mediated gene editing and haploid induction revealed that *in vivo* haploid induction can enable gene editing in maternal haploids in any background, without mixing the paternal genome (commonly referred to as “HI-edit” or “IMGE”) (Kelliher et al., 2019; Wang et al., 2019). This is promising for future genetic improvement of crops. However, the “HI-edit using CRISPR/Cas9 and interspecific hybridization has a lower editing efficiency in wheat than that in maize (Kelliher et al., 2019; Wang et al., 2019). Because our study demonstrates the feasibility of *in vivo* haploid induction in wheat, it also provides a putative platform for wheat haploid gene editing with CRISPR/Cas9. Considering the conservation of gene sequence and the capacity for haploid induction among maize, rice and wheat, this method could potentially be extended to a wider variety of crop species.

## Methods

### Knockout of homologous genes in wheat

The full-length amino acid sequence of *ZmPLA1*/*MTL*/*NLD* was used to perform a BLAST search for the homologous genes in crop species. Amino acid sequences of homologues were downloaded (http://www.gramene.com) and used for cluster alignment followed by a phylogenic analysis with the neighbor-joining method using the software MEGA X with default parameter (Knyaz et al., 2018). A guide RNA sequence was then designed targeting the homologous gene and then cloned into vector pBUEB411 (Liu et al., 2017b; Xing et al., 2014). The hexaploid wheat line CB037 was used for transformation (Wang et al., 2017). Young embryos were sampled and used for *A. tumefaciens*-mediated transformation. Positive calluses were screened using bialaphos. Transgenic T_0_ plants were transplanted in the green house and verified by sequencing of the target site. Plants with insertion/deletion mutations that lead to frame shifts and complete loss of function were chosen to produce the T_1_ generation. Genotyping was confirmed by sequencing in the T_1_ plants by sequencing were performed again.

### Haploid plant identification and verification

Putative haploids were screened in the T_1_ generation. In maize, haploids have shorter plant height, narrower leaves, smaller anther and are sterile. Thus, wheat plants with the above-mentioned phenotypes were selected as putative haploids. Subsequently, ploidy of putative haploids was verified using flow cytometry (Liu et al., 2017a). Young leaves of both hexaploids (controls) and putative haploids were sampled and chopped with a razor blade in buffer (Schutte et al., 1985). Nuclei were extracted and stained with a fluorescent dye. The signal peak of standard hexaploids (controls) was set to 100, these putative haploids with signal peak at 50 were considered real haploids. HIR was calculated as follows: HIR (%) = (number of haploid plants/total number of plants derived from same individual) × 100%. For each ear, the SSR was calculated as follows: SSR (%) = (number of viable seeds/total number of embryo sacs on the ear) × 100%.

### Pollen viability analysis

A fresh FDA staining buffer was made by mixing 100 μl FDA stock solution (0.5% in acetone) with 4.9 ml sucrose solution (Widholm, 1972). Fresh pollen (20 mg) was sampled from wildtype and mutant lines in the greenhouse and mixed immediately with the FDA working solution and left in dark environment for 1 h without covering to guarantee an abundant supply of oxygen for the esterase reaction. Three biological replicates were performed. Fluorescent was observed and photographed by a fluorescence microscope (Nikon, Ci-S-FL) with an excitation wavelength of 485 nm. Pollens that exhibited strong green fluorescence from the cytosol was considered to be highly viable pollen, pollen that exhibited weak fluorescence was considered as low viability pollen, and pollen with no detectable staining was considered dead.

### Subcellular co-localization

RNA from mature pollen of wildtype CB037 was extracted using TRIzol reagent. Reverse transcription was performed using oligo (dT) primers to obtain full-length cDNA. The coding sequence of *TaPLA*-A, *TaPLA*-B and *TaPLA*-D without termination codons (TGA) were cloned into a vector 1305-EGFP driven by the 35S promotor. Sequencing-validated constructs of 35S::*TaPLA*-A-EGFP, 35S:: *TaPLA*-B-EGFP and 35S:: *TaPLA*-D-EGFP and ER marker were used for *A. tumefaciens*-mediated transformation into tobacco epidermal cells. After culturing at 21°C for 48 hours, EGFP signals were observed and imaged using a confocal laser-scanning microscope (Zeiss LSM 710). As a control, empty 1305-EGFP vector was also transformed.

## Author contributions

S.C. and C.L. conceive and designed the project. C.L., Y.Z. constructed plasmid. X.Q., Y.Z., M.C., Z.L., C.C., X.T., J.L., D.W., Y.W., M.L. and W.L. planted transgenic plants in greenhouse and performed haploid identification and verification, phenotype investigations. Y.Z., C.L., X.Q., M.L., and W.L. performed data analysis. C.L., Y.Z., X.Q. M. X. and S.C. wrote the paper.

## Acknowledgments

We thank Prof. Xingguo Ye of Institute of crop science, Chinese academy of agricultural sciences, for his help in wheat transformation. We thank Prof. Pu Wang of College of Agronomy and Biotechnology, China Agricultural University, for providing greenhouse. We thank Dr. Qiguo Yu of Rutgers, The State University of New Jersey, for reading the manuscript. We thank Prof. Jingrui Dai, Prof. Qixin Sun and Prof. Zhongfu Ni for their valuable suggestions. This research was supported by the National Key Research and Development Program of China (2016YFD0101200) the Modern Maize Industry Technology System (CARS-02-04) and China Postdoctoral Science Foundation (2018M631634).

---

**Supplemental Figure 1.**
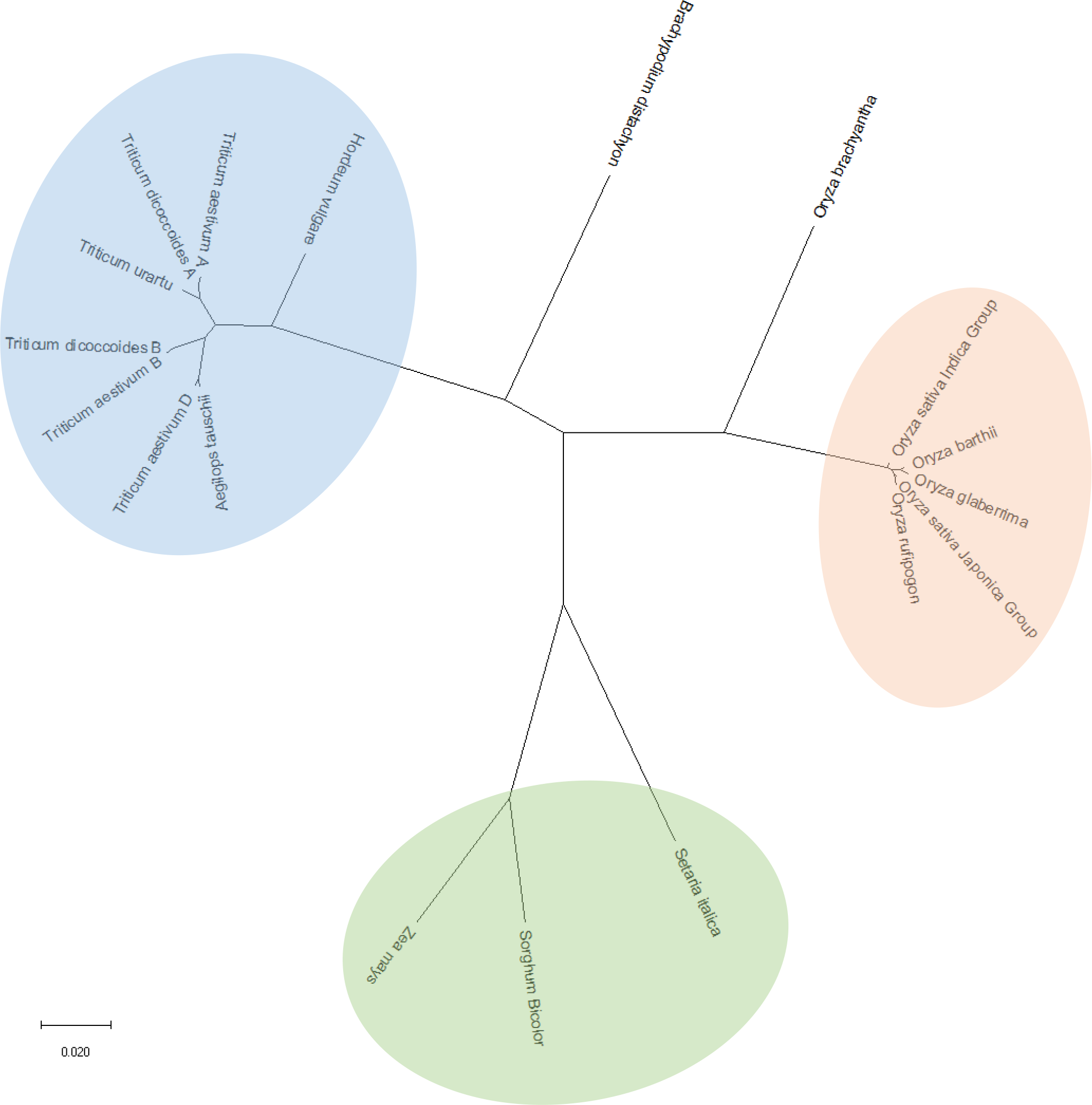
Phylogenetic analysis of *MTL*/*ZmPLA1*/*NLD* homologues among main crop species. The green background indicates phylogenetic branch containing maize, sorghum and millet; blue background indicates wheat relative species; orange background indicates rice relative species. The scale bar indicates the proportion of the sites changing along each branch.

**Supplemental Figure 2.**
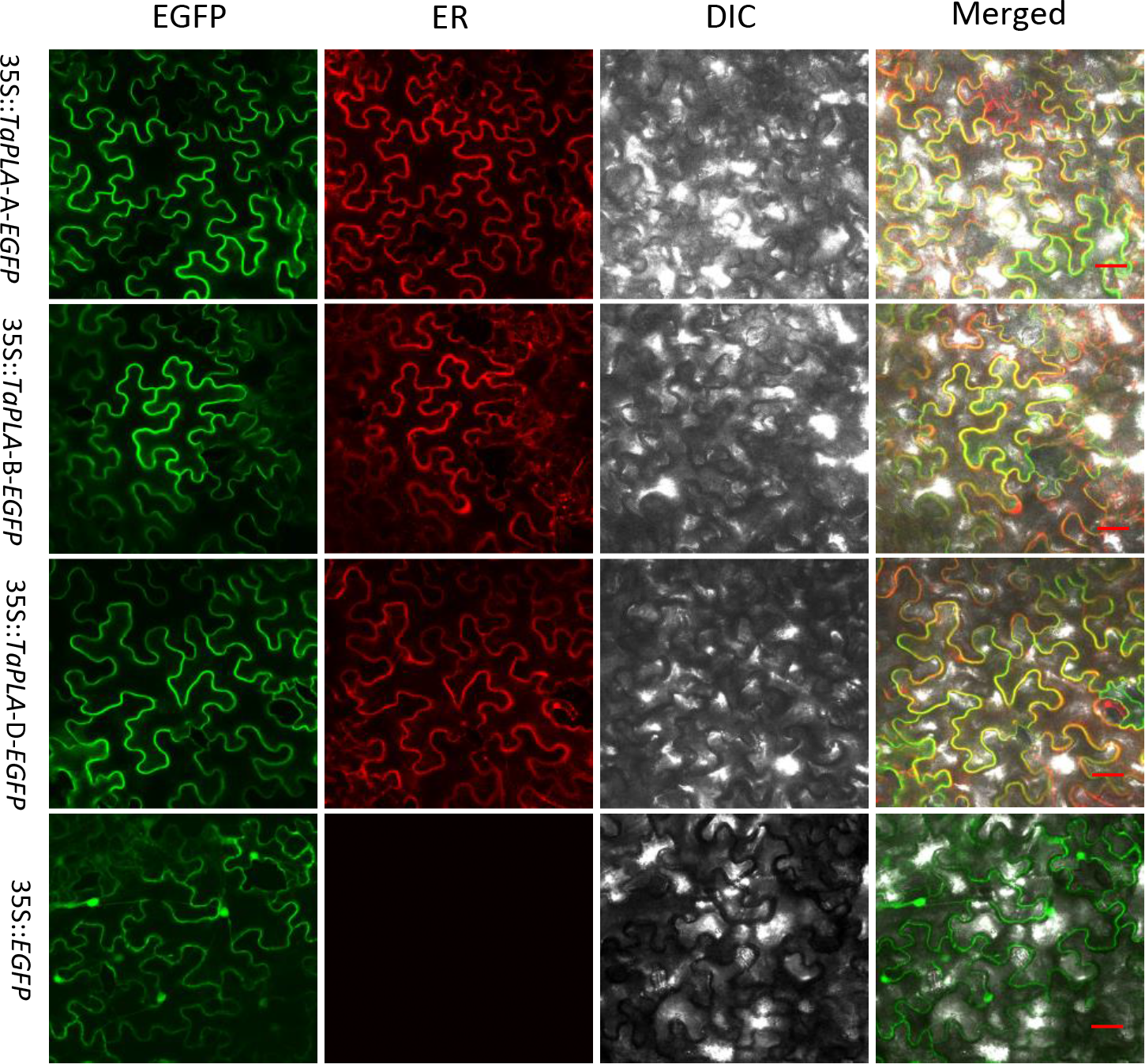
Subcellular localization of *TaPLAs* in tobacco epidermal cells. Confocal scanning (GFP; outside left), ER-mCherry signal (inside left), differential interference contrast (DIC; inside right), and merged (outside right) micrographs of tobacco epidermal cells transformed with a 35S::*TaPLA-* A-EGFP (row 1), 35S::*TaPLA-*B-EGFP (row 2), 35S::*TaPLA-* D-EGFP (row 3) and 35S::EGFP control (row 4). Scale bars = 100 μm.

**Supplemental Table 1.**
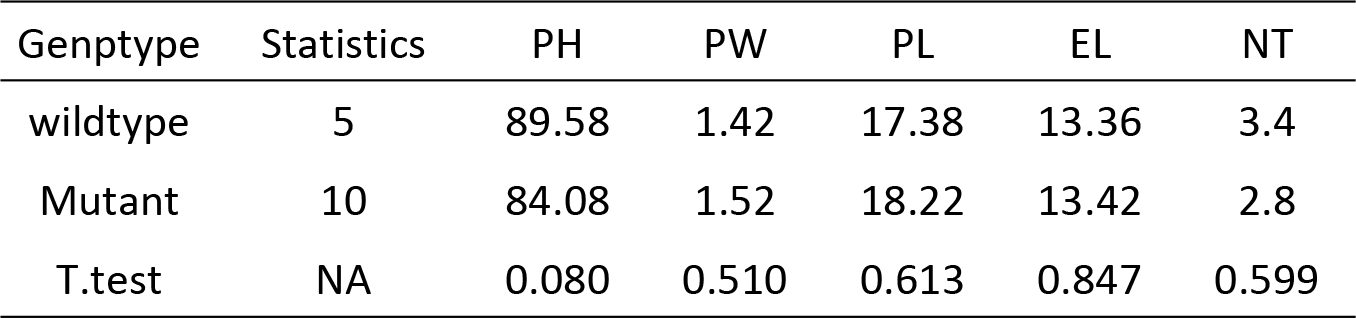
Phenotypic differences between wildtype and mutant plants. Both wildtype and double mutants are in CB037 background. PH, plant height; LW, leaf width of the flag leaf from stem; LL, leaf length of the flag leaf from stem; EL, stem ear length; NT, the number of tillers; NA, Not Applicable. Each phenotype was compared between wildtype and mutant using a heteroscedastic two-tailed Student’s *t*-test.

| Genptype | Statistics | PH | PW | PL | EL | NT |
| --- | --- | --- | --- | --- | --- | --- |
| wildtype | 5 | 89.58 | 1.42 | 17.38 | 13.36 | 3.4 |
| Mutant | 10 | 84.08 | 1.52 | 18.22 | 13.42 | 2.8 |
| T.test | NA | 0.080 | 0.510 | 0.613 | 0.847 | 0.599 |

**Supplemental Table 2.**
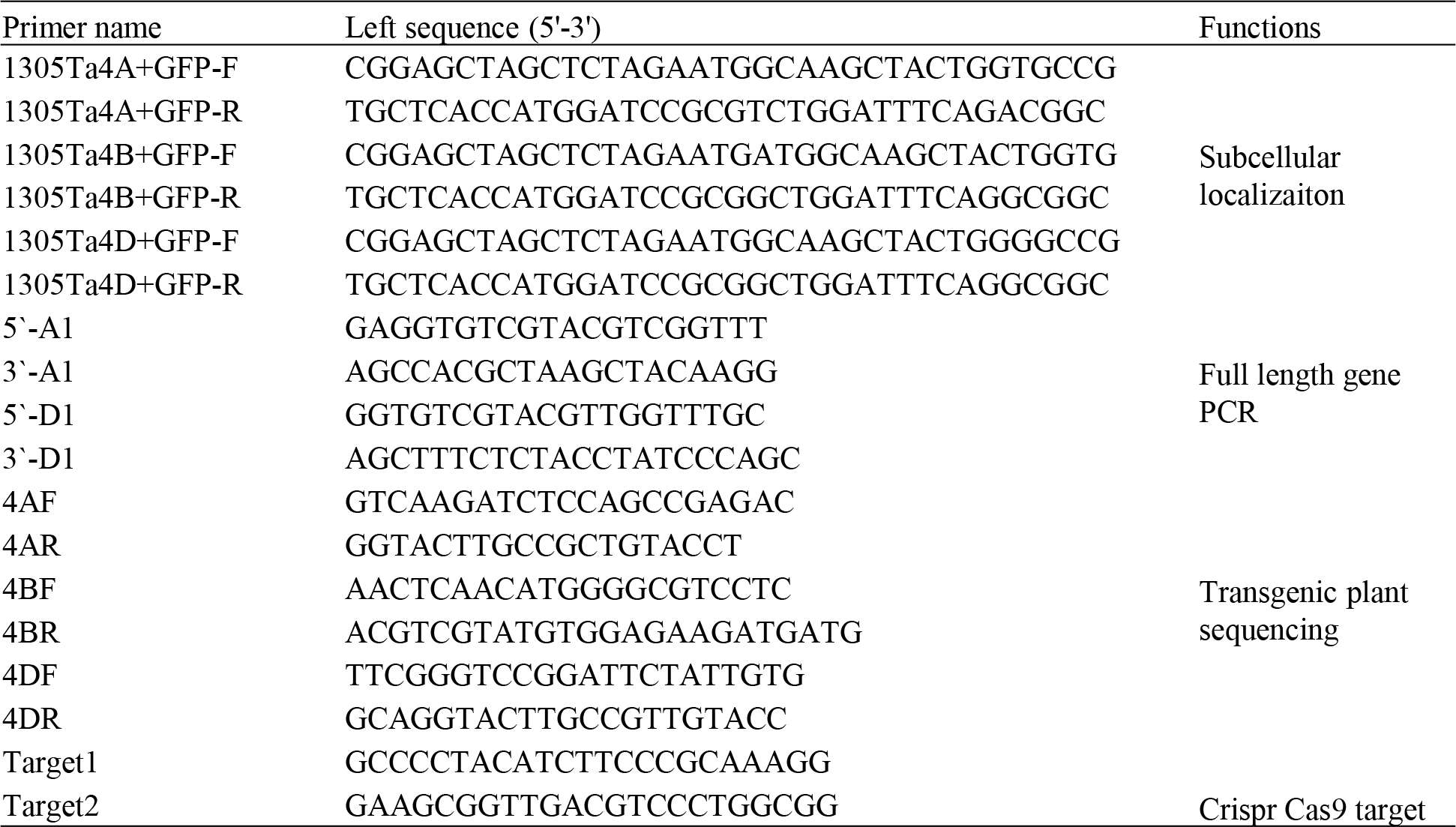
Primers used in this study.

